# Valproic acid exposure affects social visual lateralization and asymmetric gene expression in zebrafish larvae

**DOI:** 10.1101/2022.08.22.504839

**Authors:** Andrea Messina, Valeria Anna Sovrano, Greta Baratti, Alessia Musa, Alice Adiletta, Paola Sgadò

**Author notes:** Correspondence to: Paola Sgadò, PhD, Valeria Anna Sovrano, PhD. These authors contributed equally (Andrea Messina, Valeria Anna Sovrano, Greta Baratti).

## Abstract

Cerebral asymmetry is critical for typical brain function and development, at the same time altered brain lateralization seems to be associated with neuropsychiatric disorders. In autism spectrum disorders (ASD) studies have suggested reduced functional and structural cerebral asymmetry, reporting changes in asymmetric activation of brain structures involved in language and social processing, and increased prevalence of left-handedness. Zebrafish are increasingly emerging as model species to study brain lateralization, using asymmetric development of the habenula, a phylogenetically old brain structure associated with social and emotional processing, to investigate the relationship between brain asymmetry and social behavior. We exposed 5-hour post-fertilization zebrafish embryos to valproic acid (VPA), a compound used to model the core signs of ASD in many vertebrate species, and assessed social interaction, visual lateralization and gene expression in the thalamus and the telencephalon. VPA-exposed zebrafish exhibit social deficits and a deconstruction of social visual laterality to the mirror. We also observe changes in the asymmetric expression of the epithalamic marker *leftover* and in telencephalic gene expression in adult zebrafish. Our data indicate that VPA exposure neutralizes the animals’ visual field bias, with a complete loss of the left-eye use bias in front of their own mirror image, and alters brain asymmetric gene expression, opening new perspectives to investigate brain lateralization and its link to atypical social cognitive development, considering zebrafish as an animal model for the autistic syndrome.

## Introduction

Functional and structural lateralization has been documented in humans^1–3^ and many other vertebrate species^4,5^, including fish^6,7^. Human and animal model studies suggest a general pattern of specialization of the two hemispheres^4,8^: the right hemisphere seems to be specialized for social and emotional processing and response to danger and novelty^9–13^, while the left hemisphere attends to categorization and attention, and controls the fine motor skills that underly language dominance and the population-level right-handedness in humans and other animals^14,15^. Hemispheric dominance and functional lateralization seem to be critical for typical cognitive development^16^. Loss of cerebral lateralization (either weaker or absent asymmetry) underlies poorer cognitive abilities, and in some cases is associated with brain disorders, including autism^17^. Autism Spectrum disorder (ASD) comprise a heterogeneous group of conditions characterized by atypical social interaction and communication, restricted interests and repetitive behaviour, and sensory processing abnormalities. A large body of literature suggests alterations in hemispheric functional asymmetry associated with ASD, emerging since early development^18^. Deficits in language processing^19–22^ and abnormal hemispheric activation in response to speech^23^ have been consistently described in ASD. Interestingly, atypical prevalence of handedness have also been observed in individuals with ASD^24^. In vertebrates with front-facing eyes and binocular vision, such as humans, lateralized processing can be documented and measured through the observation of visual field biases, as, for example, the strong left visual field bias demonstrated in humans in face detection^25,26^ and emotional processing^see for a review 27^. In addition to aberrant lateralized responses to speech and in language processing, lack of left visual field bias in face and emotional processing have been extensively reported in ASD^28–30^, suggesting altered functional and structural lateralization already at early developmental stages^31^. On the same line, neurophysiological and neuroimaging studies have indicated altered patterns of lateralized activation in cortical brain areas associated with configurational information and categorization of faces (e.g., fusiform face area)^32–36^.

In recent years, studies in vertebrate species displaying functional lateralization have significantly contributed to widening the knowledge about the mechanisms underlying brain asymmetry and its role in cognitive functions^4^. Zebrafish are increasingly emerging among the key model species to study functional and anatomical aspects of brain asymmetry^7^. Similar to the visual field biases measured in humans, thanks to the bilateral positioning of the eyes and an almost complete decussation of the optic chiasm, the perception and processing of stimuli in zebrafish can be inferred, already at early developmental stages, on the basis of the simple observation of eye dominance during spontaneous behaviour^37^.

Studies in zebrafish larvae suggest that social stimuli are processed by the right hemisphere, as revealed by a left visual field bias the larvae show while observing their image reflected in a mirror^38^. In addition to behavioural lateralization, the asymmetric development of the zebrafish epithalamus have been used as a model to study the relationship between brain asymmetry and behavior^39,40^. During development of the zebrafish epithalamic region, the parapineal organ is located on the left side in most of the embryos, asymmetrically influencing the development of the habenular nuclei. As a consequence the two dorsal habenula nuclei show differences in size, connectivity^41–44^ and in the expression of the *kctd12*.*1/leftover* gene, which is typically asymmetrically distributed, highly expressed in the left habenula compared to the right one^43^. Two other members of the same gene family, *ktcd12*.*2/right-on* and *ktcd8/dexter* have an opposite expression pattern^44^.

Given the fundamental contribution of cerebral asymmetry in brain organization and development, and its association with neuropsychiatric diseases, including autism, aim of this study was to investigate brain lateralization in an animal model of ASD based on embryonic administration of valproic acid (VPA), a compound used to model the core signs of ASD in many vertebrate species^45–47^. Five-hour-post-fertilization (5 hpf) zebrafish larvae were exposed to one micromolar VPA for 24 and 48 hours and tested, at three- and four-weeks post-fertilization, for the social responses to their reflected image (mirror test) and to the conspecifics (social preference), respectively. At three-months-post-fertilization VPA-exposed zebrafish were assessed for asymmetric gene expression in the thalamus and in the telencephalon. VPA exposure induces social interaction deficits and a deconstruction of social visual laterality, and alters biological pathways underlying brain lateralization.

## Results

### Mirror test

One hundred twenty-four larvae of the AB strain, treated with vehicle (39), 1μM VPA for 24 (46) or 48 hours (39), were subjected to the mirror test (Figure 1A) to assess the effect of treatment on the left visual field bias. The left visual field bias was expressed as the ratio of left eye use when the fish were observing their reflection close to the mirror. The analysis of variance indicated a significant effect of treatment on the left visual field index (Figure 1B; F_(2,121)_ = 27.76, p <0.0001), with a remarkable reduction, at the population level, of the use of the left eye during the test in both the VPA 24 and 48 hours treatment group (Figure 1B; pairwise comparisons: CTRL vs VPA 24h t_(121)_= 5.980, p < 0.0001; CTRL vs VPA 48h t_(121)_= 6.908, p < 0.0001; VPA 24h vs VPA 48h t_(121)_= 1.206, p = 0.4518). In accordance with previous results, the vehicle-treated larvae displayed a stable bias for left eye use (Figure 1B; one-sample t-test: CTRL t_(38)_ = 10.317, p < 0.0001; group mean: CTRL 0.5960 [95% C.I. 0.5772 – 0.6148]), while both VPA treatment groups showed no preferential visual field use during the test (Figure 2B; one-sample t-test: VPA24h t_(45)_ = 1.4861, p = 0.1442; VPA48h t_(38)_ = -0.2105, p = 0.8344; group mean: VPA24h 0.5143 [95% C.I. 0.4949 – 0.5337]; VPA48h 0.4978 [95% C.I. 0.4771 – 0.5186]). This data suggests that VPA treatment impairs the behavioural lateralization shown by the larvae in response to their reflected image, as visual cues representing a conspecific.

**Figure 1.**
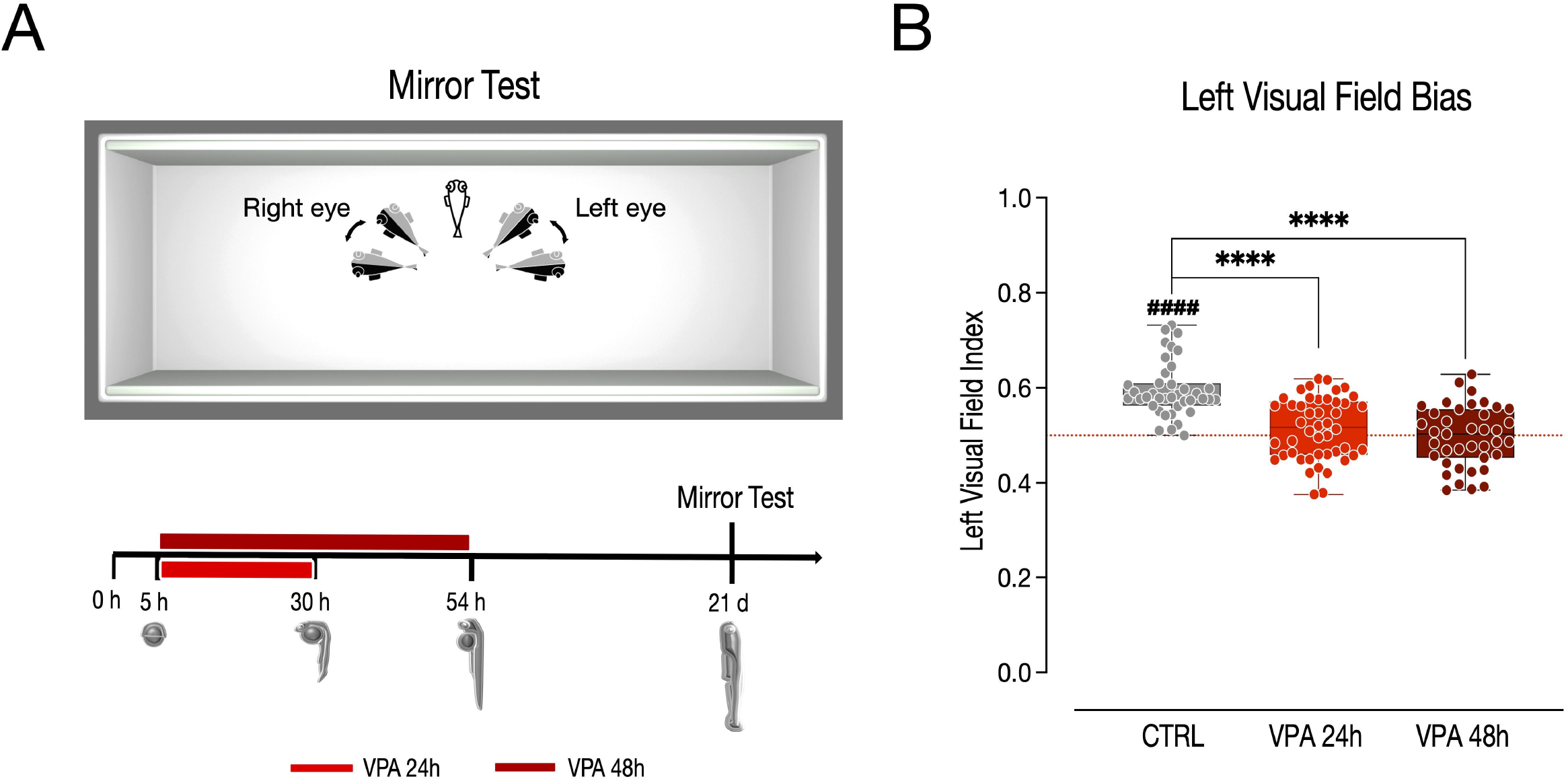
Mirror test. (A) Top, apparatus used for the mirror test, showing the position of the mirrors and the angles of viewing that defined monocular vision with the right or left eye. Data were discarded when the fish was perpendicular to the mirror (binocular stimulation, transparent fish) or when it formed an angle larger than 90º with respect to the closest mirror. Bottom, scheme of the experimental timeline, VPA treatment begin at 5 hpf and last for 24 or 48 hours. The mirror test starts at 21 dpf. (B) Box and whisker plot (median, min to max) showing the left visual field index. The number sign (#) indicate significant departures of the left visual field index from chance level (0.5), marked by the red line. ^####^p < 0.0001; ****p < 0.0001.

**Figure 2.**
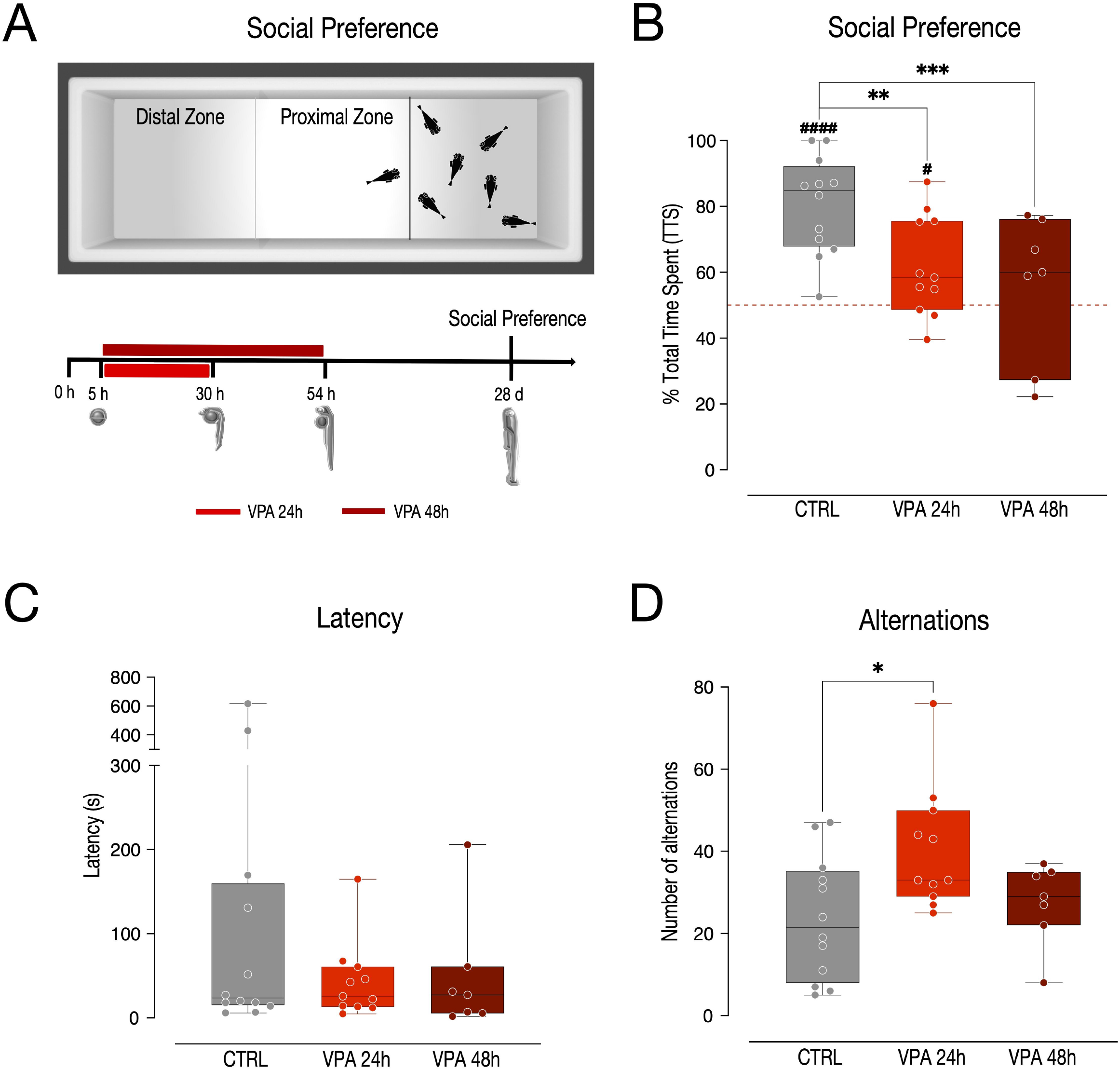
Social preference test. (A) Top, apparatus used for the social preference test, showing the conspecifics chamber and the two areas, the proximal and distal zone. Bottom, scheme of the experimental timeline, VPA treatment begin at 5 hpf and last for 24 or 48 hours. The social preference test starts at 28 dpf. (B, C, D) Box and whisker plot (median, min to max) showing (B) the % of time spent in the proximal zone (social preference index), (C) the latency to change zone and (D) the number of alternations between proximal and distal zones. The number sign (#) indicate significant departures of the social preference index from chance level (50%), marked by the red line. #p < 0.05; ^####^p < 0.0001; *p < 0.05; **p < 0.01; ***p < 0.001.

### Social interaction test

To assess the effect of VPA on a more general sociability test, an independent group of larvae was tested in the social interaction test. Thirty larvae of the AB strain treated with vehicle (12), 1μM VPA for 24 hrs (11) or 48 hrs (7) underwent the social preference tests at four weeks of age (Figure 2A). To account for potential habituation of the fish to the environment, the analysis of variance also evaluated the effect of time (three and five minutes, repeated measures) on the social preference and its interaction with VPA treatment. The results show no effect of time (F_(1,54)_ = 0.283, p = 0.597) nor of the interaction between time and treatment (F_(2,54)_ = 0.107, p = 0.899) but a significant effect of treatment (F_(2,54)_ = 11.813, p < 0.0001) on the social preference index. Twenty-four and 48 hours of treatment with VPA significantly decreased the preference of the fish to spend time in the chamber close to the conspecifics (Figure 2B; pairwise comparisons: CTRL vs VPA 24h t_(54)_= 3.791, p = 0.0011; CTRL vs VPA 48h t_(54)_= 4.352, p = 0.0002; VPA 24h vs VPA 48h t_(54)_= 1.008, p = 0.5750). A very strong bias to remain in the compartment of the chamber adjacent to the conspecifics was detected in fish treated with vehicle, while the group of animals treated with VPA for 48 hours did not show any preference for the proximal chamber (Figure 2B; one-sample t-test: CTRL t_(11)_ = 7.083, p < 0.0001; VPA24h t_(10)_ = 2.592, p = 0.0269; VPA48h t_(6)_ = 0.659, p = 0.5343; group mean: CTRL 80.43% [95% C.I. 70.97% - 89.88%]; VPA24h 61.93% [95% C.I. 51.67% - 72.18%]; VPA48h 55.54% [95% C.I. 34.96% - 76.12%]). To assess potential confounds on the social preference related to motor activity, the latency to the first change of zone and the number of alternations between the two zones during the social preference test were also evaluated. The statistical analysis showed no significant effect of treatment on the latency (Figure 2C; F_(2,27)_ = 1.320, p = 0.2839), however a clear effect of treatment on the number of spontaneous alternations emerged (Figure 2D; F_(2,27)_ = 4.385, p = 0.0224). Larvae treated with 1 μM VPA for 24 hours, but not for 48 hours, displayed a significant increase in the number of alternations between the proximal and the distal zone (Figure 2D; pairwise comparisons: CTRL vs VPA 24h t_(27)_ = -2.881, p = 0.0203; CTRL vs VPA 48h t_(27)_ = -0.586, p = 0.8288; VPA 24h vs VPA 48h t_(27)_ = 1.911, p = 0.1550).

### Asymmetric gene expression analysis

To investigate whether the functional lateralization disturbances mediated by VPA in the mirror test were also associated with alterations in brain asymmetry, the left and right thalami of three-month-post-fertilization (mpf) zebrafish were micro-dissected and the expression levels of the thalamic markers *kctd12*.*1/leftover* (*lov*), *ktcd8/dexter* (*dex*) and *ktcd12*.*2/right-on* (*ron*) evaluated (Figure 3A; n = 6 animals per treatment group, 6 independent experiments). Considering that behavioural analyses did not reveal substantial differences between zebrafish exposed to same concentrations of VPA for 24 hours or for 48 hours, gene expression was analyzed only in those larvae treated for 24 hours. To assess the effect of treatment on the asymmetric expression of these genes, we fitted a linear mixed model, including treatment, brain side and transcript type as fixed factors and the experimental unit (experiment) as random factor. We compared a model with random-intercepts-only to one with random slopes and intercepts and found that the second model fitted the data significantly better. The statistical analysis indicated a significant difference in asymmetric gene expression in the treatment groups (interaction treatment * brain side F_(1,55)_ = 13.8159, p = 0.0005). We also found a significant difference in the expression levels of the three transcripts analyzed (main effect of transcript F_(2,55)_ = 1332.2072, p < 0.0001) and no other significant main effects or interactions (main effect of treatment F_(1,55)_ = 0.2256, p = 0.6367; main effect of brain side F_(1,55)_ = 0.6442, p = 0.4257; treatment * transcript interaction F_(2,55)_ = 2.6251, p = 0.0815; brain side * transcript interaction F_(2,55)_ = 2.6440, p = 0.0801; treatment * brain side * transcript interaction F_(2,55)_ = 1.7248, p = 0.1877). The pairwise comparison between the levels of expression in the two hemispheres of each treatment groups revealed an asymmetric expression of *kctd12*.*1/leftover* in the control samples, with a higher expression on the left hemisphere compared to right one, in line with previous reports^44,48,49^. This difference was however absent in zebrafish treated with VPA (Figure 3B and Table 1; *leftover* CTRL right – left: t_(55)_ = -2.930, p = 0.0049; *leftover* VPA 24 hrs right – left: t_(55)_ = 1.195, p = 0.2374). No changes in the expression of *ktcd12*.*2/right-on* was detected between the two hemispheres in the treatment groups (Figure 3B and Table 1; *right-on* CTRL right – left: t_(55)_ = 0.980, p = 0.3313; *right-on* VPA 24 hrs right – left: t_(55)_ = 1.871, p = 0.0667), nor in the expression of *ktcd8/dexter* in the control samples, while in VPA treated samples *ktcd8/dexter* was expressed at lower levels in the left hemisphere compared to the right one (Figure 3B and Table 1; *dexter* CTRL right – left: t_(55)_ = -1.619, p = 0.1112; *dexter* VPA 24 hrs right – left: t_(55)_ = 2.470, p = 0.0166; see Table 1 for a summary of all pairwise comparisons). The analysis of the Lateralization index also indicate, as previously reported at different developmental stages^44,48,49^, that *kctd12*.*1/leftover* expression was asymmetric, predominant on the left side of the thalamus in the control group at this stage, as shown by the significant departure from chance of the lateralization index (Figure 3C and Table 2; one sample t-test of left lateralization index *leftover* CTRL t_(5)_ = 11.73, p < 0.0001). This lateralized expression was however absent in VPA treated animals (Figure 3C and Table 2; one sample t-test of left lateralization index *leftover* VPA 24h t_(5)_ = 1.235, p = 0.2716). Differently from previous reports describing the asymmetric distribution of *ktcd8/dexter* and *ktcd12*.*2/right-on* primarily on the right hemisphere during development^44,48,49^, these thalamic markers were not clearly preferentially distributed in one of the hemispheres of the adult thalamus in any of the treatment groups (Figure 3C and Table 2; one sample t-test of left lateralization index *dexter* CTRL t_(5)_ = 2.216, p = 0.0775; *dexter* VPA 24h t_(5)_ = 2.326, p = 0.0675; *right-on* CTRL t_(5)_ = 0.7083, p = 0.5104; *right-on* VPA 24h t_(5)_ = 1.587, p = 0.1773; see Table 2 for a summary of all one-sample t-test results).

**Figure 3.**
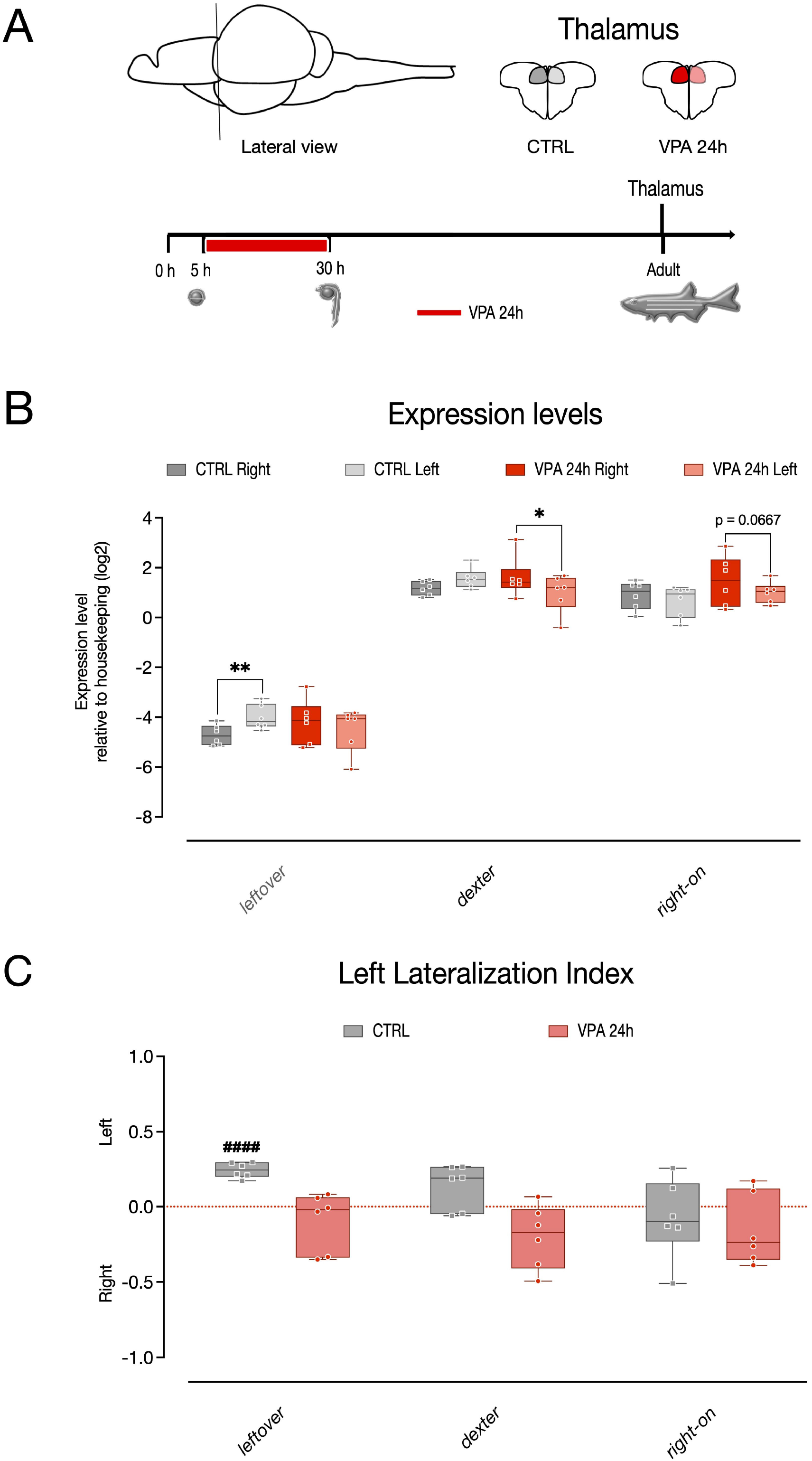
Gene expression in the left and right thalamus of zebrafish. (A) Top, schematic representation of thalamic regions selected for the analyses. Bottom, scheme of the experimental timeline, VPA treatment begin at 5 hpf and last for 24 hours. Gene expression is analyzed in adult zebrafish. (B) Box and whisker plot (median, min to max) of relative expression (dCt, log2) values for each treatment group for *kctd12*.*1/lov, kctd8/dex* and *kctd12*.*2/ron* in the left and right thalamus of zebrafish. (C) Left lateralization index for *kctd12*.*1/lov, kctd8/ dex* and *kctd12*.*2/ron*. The number sign (#) indicate significant departures of the left lateralization index from chance level (0.5), marked by the red line. ^####^p < 0.0001; *p < 0.05; **p < 0.01.

To assess brain asymmetry also in the telencephalon, expression of pallial genes previously shown to be asymmetrically distributed in adult zebrafish^49^ was examined in VPA-exposed zebrafish. The left and right telencephala (including the pallium and the subpallium) were dissected and then the expression of *arrb2, fez1, gap43, nipa1, nipa2 and robo1* evaluated in the two hemispheres of 3 mpf zebrafish subjected to same treatment paradigm: 1μM VPA at 5 hpf for 24 hrs (Figure 4A; n = 4 animal per treatment group, 4 independent experiments). To assess the effect of treatment, brain side and transcript type on the expression levels, we again fitted a linear mixed model, including the experiment (experimental unit) as a random factor. Using a random slopes and intercepts model, the analysis revealed a significant asymmetric gene expression in the treatment groups (Figure 4B; interaction treatment * brain side F_(1,69)_ = 8.5682, p = 0.0046) as well as a difference in the expression levels of the transcripts analyzed (main effect of transcript F_(5,69)_ = 1125.7494, p < 0.0001) and an overall difference in the expression levels between hemispheres (main effect of brain side F_(1,69)_ = 17.3302, p = 0.0001). No other significant main effects or interactions emerged (main effect of treatment F_(1,69)_ = 0.0695, p = 0.7928; transcript * brain side interaction F_(5,69)_ = 1.0046, p = 0.4217; treatment * transcript interaction F_(5,69)_ = 0.1690, p = 0.9732; treatment * brain side * transcript interaction F_(5,66)_ = 0.4750, p = 0.7937). The pairwise comparison between the expression levels of the two hemispheres in the treatment groups indicated no difference in the asymmetric expression in the control group of all genes analyzed, differently from what previously described^49^, while a significant difference between hemispheres was observed in the VPA treatment group for the gene *gap43, nipa2* and *robo1* that showed increased expression in the left hemisphere (Figure 4B and Table 3; *gap43* VPA 24 hrs right – left: t_(69)_ = -2.279, p = 0.0258; *nipa2* VPA 24 hrs right – left: t_(69)_ = -3.152, p = 0.0024; *robo1* VPA 24 hrs right – left: t_(69)_ = -3.377, p = 0.0012; see Table 3 for a summary of pairwise comparison results). Differently from previous reports^49^ the left lateralization index for all the genes analyzed was not suggestive of lateralization in their distribution in the control groups, however in some cases the expression was asymmetrically distributed in the VPA-treated group, as for example in the case of *arrb2, fez* and *nipa2*, for which significant departure from chance levels of the left lateralization indexes was observed (Figure 4C and Table 4; one sample t-test of left lateralization index *arrb2* VPA 24h t_(3)_ = 5.336, p = 0.0129; *fez* VPA 24h t_(3)_ = 3.272, p = 0.0336; *nipa2* VPA 24h t_(3)_ = 5.818, p = 0.010; see Table 4 for a summary of all one-sample t-test results).

**Figure 4.**
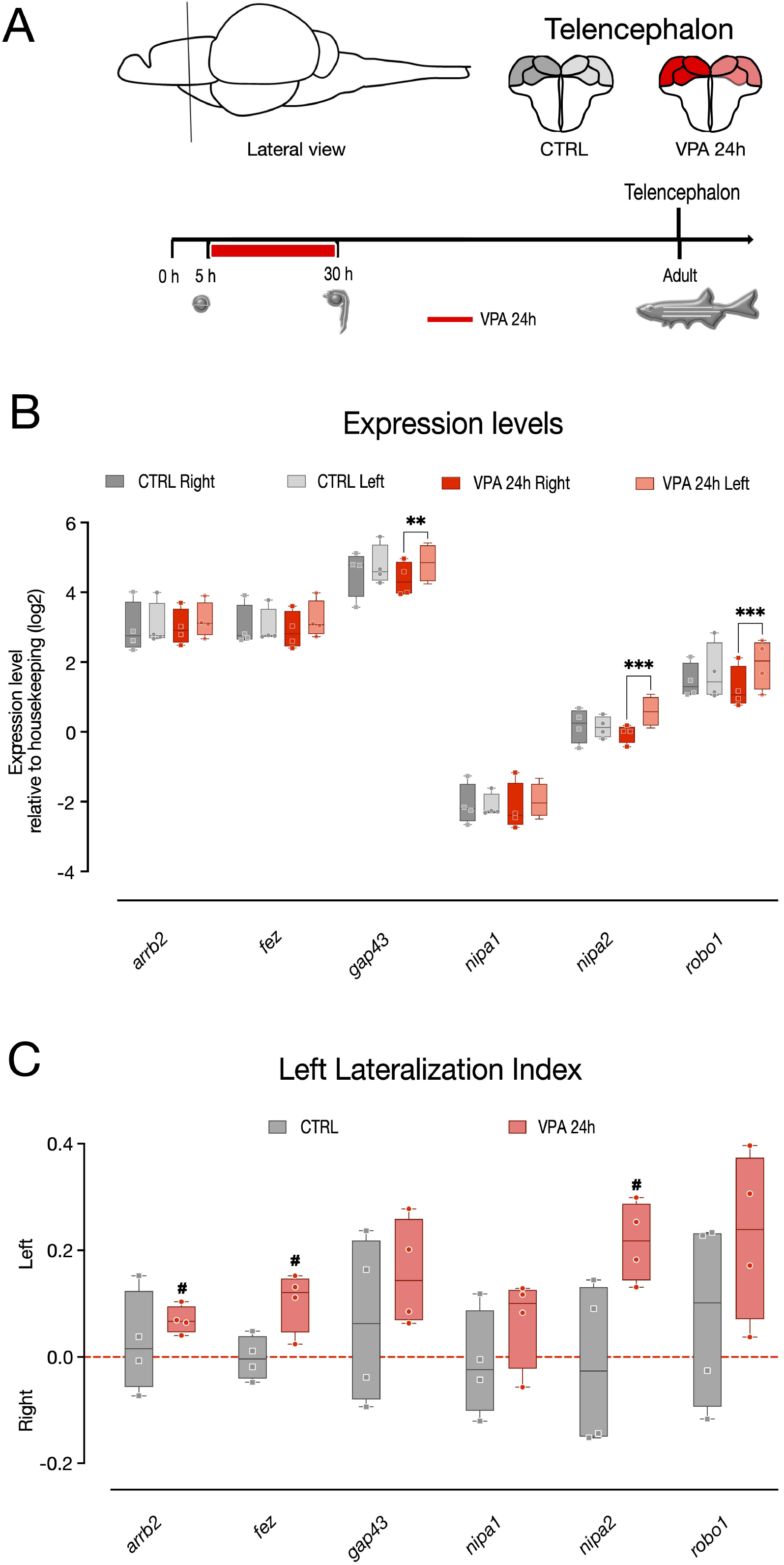
Gene expression in the left and right telencephalon of zebrafish. (A) Top, schematic representation of telencephalic (pallial and subpallial) regions selected for the analyses. Bottom, scheme of the experimental timeline, VPA treatment begin at 5 hpf and last for 24 hours. Gene expression is analyzed in adult zebrafish. (B) Box and whisker plot (median, min to max) of relative expression (dCt, log2) values for each treatment group for *arrb2, fez, gap43, nipa1, nipa2* and *robo1* in the left and right telencephalon of zebrafish. (C) Left lateralization index for *arrb2, fez, gap43, nipa1, nipa2* and *robo1*. The number sign (#) indicate significant departures of the left lateralization index from chance level (0.5), marked by the red line. ^#^p < 0.05; **p < 0.01; ***p < 0.001.

## Discussion

Research focusing on language^50–53^ and face processing^28–31^ have demonstrated changes in functional lateralization in autistic children of young age. Although cerebral asymmetry is recognized to be critical for cognitive development^37^ the role of brain lateralization in atypical social development is still unclear. Animal models are fundamental tools to investigate neurobiological mechanisms associated with human disorders, providing mechanistic insights into evolutionarily conserved functions. Here, we have investigated behavioral and biological lateralization in a zebrafish model of ASD based on embryonic administration of VPA. A single embryonic exposure to VPA induces neuroanatomical and behavioural changes that resemble the core signs of ASD in many vertebrate species^54,55^. Previous studies in zebrafish have established a detrimental effect of VPA on social behaviour^56–58^ at different dosages. A previous report^59^ has identified the lowest dose of VPA (1μM) that minimizes the toxic effect on zebrafish embryo survival, maintaining its effect on the expression of several neurodevelopmental genes previously known to be affected by VPA^60,61^. However, the effect of lower doses of VPA on social behaviour was not assessed. This study shows that one micromolar VPA dose not only induces social interaction deficits but also affects the left visual bias in a social recognition task, based on the animals’ response to their reflection in a mirror. Moreover, the gene expression data suggests that the deficits in behavioural lateralization were accompanied, in three-month-old zebrafish, by changes in asymmetrically distributed biological pathways linked to developmental brain lateralization, as indicated by the changes in the thalamic marker *kctd12*.*1/leftover*. This data demonstrates that exposure to one micromolar VPA is sufficient to induce social preference deficits and to neutralize both the left visual bias and the asymmetric expression of *kctd12*.*1/leftover*. Given the importance of cerebral asymmetry for brain development and function, this work may open new perspectives on the study of brain lateralization and its link to typical and atypical social development in animal models of the human disorders.

Similar to the social affiliative responses shown by other vertebrate species towards visual cues in the face region^62–64^, zebrafish larvae exhibit spontaneous approach responses to their reflected image, reacting to the mirror image as to a social stimulus representing a conspecific^65,66^. Interestingly, zebrafish larvae observing their mirror image also display a stable left visual bias^38^ that is shared with other fish species^67–70^ and reaches a maximum at the same stage when social interaction begin to emerge (i.e. the third postnatal week)^71^. This study analyzes the effect of VPA exposure on this social behavioural lateralization, revealing a dramatic effect of VPA on the left visual field bias displayed by zebrafish larvae towards their reflected image. Both the 24- and 48-hours VPA treatment groups showed severe impairment of the left visual bias, reducing their laterality index to chance levels, indication of a “symmetrization” of the response to the social cues. VPA-treated zebrafish larvae thus fail to exhibit the typical lateralized response underlying the right hemisphere dominance for social recognition described in many vertebrate species, including humans^see for a review 37^.

The earliest sign of brain asymmetry in zebrafish is the activation of the Nodal pathway starting at about 18 hpf in the left epithalamus^72^. Shortly after the activation of the Nodal pathway, the parapineal appears on the left side of the pineal anlage and induces the left lateralization of the habenular nuclei^43,73^. The habenular nuclei show left-right differences in their size, their neuropil density and innervation pattern and the expression of the *kctd12*.*1/leftover* gene as early as 2 dpf^43^. Expression of the *kctd12*.*1/leftover* gene is originally asymmetric, being highly expressed on the (left) side closely opposed to the parapineal, and differently from other components of the Nodal pathway, continue to be highly expressed in the left epithalamus into adulthood^43^. Interestingly, in the small proportion of larvae with disrupted left-right asymmetry that develop bilateral parapineal organs, both habenular nuclei show high expression of the *kctd12*.*1/leftover* gene^43^. Wnt signaling also seems to be necessary for the communication between the parapineal organ and the left habenular nuclei to establish asymmetric activation of Nodal pathway genes in the brain^74^.

Given that the epithalamus is one the first region of the zebrafish brain found to have prominent brain asymmetry, several studies have examined the functional relevance of loss or reversals of left epithalamic asymmetry in zebrafish induced by perturbation of habenular identity or disruption of the parapineal organ using genetic, pharmacological approaches or physical ablation^75–80^. The predominant view is that a reversal of the epithalamic asymmetry induces inverted or absent behavioural lateralization, with substantial differences underlying the different zebrafish strain, artificial selection or genetic manipulation^76–79^. Our behavioural and molecular analyses are consistent with the idea that perturbations of epithalamic asymmetry affect the larvae behavioral lateralization, demonstrating that embryonic treatment with VPA alters the expression of the gene *kctd12*.*1/leftover*, increasing its expression on the right side, and suggest that VPA affects the biological mechanisms underlying brain lateralization. Interestingly, VPA exposure seems to induce a scenario similar to that seen in a small proportion of embryos in which left-right asymmetry is disrupted at early developmental stages due to abnormal midline development or to defective Nodal signaling^43^, with *kctd12*.*1/leftover* expression in the right side reaching similar levels compared to the left side (Figure 3B). As for the lack of asymmetric expression in the other thalamic markers *ktcd12*.*2/right-on* and *ktcd8/dexter* in the control zebrafish, the neuroanatomical heterogeneous nature of the adult epithalamus could account for the differences compared to what previously shown during embryonic development. Interestingly, for all epithalamic markers analyzed, VPA seem to have a similar effect, increasing their expression on the right hemisphere compared to the left one (see Figure 3C). Notably, while this study provides clear evidence of an effect of VPA on behavioural lateralization of zebrafish at three weeks and of an altered expression of *kctd12*.*1/leftover* at three months of age, the data does not provide direct support for alterations of brain asymmetry at developmental stages. Nevertheless, given that VPA treatment occurs in the first (5) hours post-fertilization, a time when the Nodal pathway starts to be active, and that VPA induces persistent changes in the brain, as demonstrated many studies analyzing the effect of embryonic exposure of VPA on adult animals^55^, it makes sense to deduce that the detrimental effect of VPA on brain lateralization may already occur at early stages.

In addition to brain asymmetry in the epithalamus, this study also analyzed the expression of pallial genes known to have a left-right asymmetric distribution in humans^81^, mice^82^ and zebrafish^49^. No asymmetric expression of the analyzed genes was detected in vehicle-treated zebrafish, while VPA treatment induced a left lateralization of the expression of *arrb2, fez* and *nipa2*. As for the lack of correspondence between the left-right distribution of these pallial markers in control animals, one potential confounds between the present study and the one from Messina et al.^49^, reporting asymmetrical expression of the same genes at the same stage, could be the inclusion in our samples of both pallial and subpallial regions. In addition, the expression of the pallial genes might also be influenced by the neuroanatomical distribution of defined cell subpopulation. Previous studies have already demonstrated an asymmetric distribution of GABAergic interneurons in the sensory cortex of mouse models for ASD^83^, including in animals exposed to VPA^84^.

Even though a clear picture on the mechanism of action of VPA on the brain developmental trajectory cannot yet be delineated, since brain asymmetry is established in the zebrafish brain at very early stages of development, before the maturation of the complex brain structures underlying social behaviour, this data seems to suggest that VPA affects brain lateralization at its initial steps, maintaining a long-term effect, eventually influencing the animals’ social cognitive abilities. It remains to be established *whether* VPA acts on both brain lateralization and social cognitive development independently and *how* brain lateralization contributes to social cognitive development.

This study shows, for the first time, that one micromolar doses of VPA are sufficient to induce social interaction deficits and to neutralize both the left visual bias in a social recognition task as well as the asymmetric expression of the epithalamic marker *kctd12*.*1/leftover*, suggesting new perspectives on the effect of VPA on brain development and proposing a new tool to investigate brain lateralization and its link to ASD in a zebrafish model.

### Limitations of the study

Despite the strong effect of VPA on the expression of the thalamic marker *kctd12*.*1/leftover*, the qPCR analyses did not detect any lateralization in the control zebrafish of the other thalamic markers, *ktcd12*.*2/right-on* and *ktcd8/dexter*. This different asymmetrically distribution in gene expression may reflect neuroanatomical differences at the levels of neuronal subpopulations compared to what previously reported in the developing epithalamus. For example the *kctd12*.*1/leftover* gene was reported to localize exclusively in dorso-lateral habenula cells^44^, while *ktcd12*.*2/right-on* is a marker for the dorso-medial cell population^48^. Moreover, given the small size of the larvae brain, asymmetric gene expression can only be assessed in three-month-old zebrafish, a considerable later stage compared to most reports.

## Acknowledgements

We thank Dr. Tommaso Pecchia for help with the experimental apparatus, Grazia Gambardella and Roberta Guidolin for administrative help and Ciro Petrone, Michela Maffei and Giampaolo Morbioli for animal facility management, welfare, and animal care.

## Authors’ contributions

AM, PS and VAS contributed to conceptualization; AM and GB contributed to methodology; AM and PS contributed to formal analysis; AM, GB, AM, AA and VAS contributed to investigation; PS contributed to writing and original draft preparation; AM, PS and VAS contributed to writing, review and editing. All authors have read and agreed to the published version of the manuscript.

## Declaration of interests

The authors declare that they have no competing interests.

## Materials and Methods

### Ethical Regulations

All husbandry and experimental procedures complied with the European Directive 2010/63/EU on the protection of animals used for scientific purposes and were approved by the Scientific Committee on Animal Health and Animal Welfare of our University and by the Ministry of Health (Protocol n. 333/2021-PR).

### Animals

Adult AB wild-type zebrafish were moved into breeding tanks overnight separated by a transparent barrier. The day after, the barrier was removed, and fish were left to breed. Embryos were collected in E3 medium (5.00 mM NaCl, 0.44 mM CaCl_2_, 0.33 mM MgSO_4_ and 0.17 mM KCl). At 5 hpf embryos were placed into 10 cm Petri dishes containing E3 medium (control) and E3 medium with 1 μM VPA (Sigma-Aldrich, P4543; Merck Life Science Srl, Milan, Italy) for 24 or 48 h. At the end of the treatment, the medium was replaced by E3 medium and zebrafish larvae were grown at 28.5°C for three-or four-weeks post-fertilization before to be subjected to behavioural studies (mirror test and social preference, respectively) or until adulthood (three months post-fertilization) before to proceed to brain microdissection.

### Mirror test

One hundred twenty-four larvae of the AB strain were observed at three weeks of age (Figure 1A). Each experimental group was composed of 39 vehicle-treated controls (CTRL), 46 and 39 larvae treated with 1μM VPA for 24 and 48 hours, respectively. Each group was observed only once. The apparatus consisted of a circular amaranth tank (diameter × height: 175 × 27 cm), surrounded by a circular black curtain fixed on a wood- and-metal frame. The mirror test apparatus was placed inside the bigger tank and was composed of white plastic walls (20 × 5 × 8 cm), with mirrors on the long walls (one per side) and lit from above (height: 100 cm) through a 24-watt fluorescent white light tube (Lumilux, Osram GmbH, D) (Figure 2A). The water was 2.5 cm deep and its temperature was maintained at 26 ± 1°C by using a 50-watt heater (NEWA Therm®, NEWA). Each larva was placed in turn in the center of the test apparatus and video-recorded from above through a webcam (LifeCam Studio, Microsoft) for 5 min. The positions of each larva were manually scored offline every 2 s, by superimposition on the computer screen of a cursor on the long axis of the body, using the video recording. Body angle was taken relative to the closest of the two mirrors. All the positions where the larva was in a central 4 mm wide area were discarded. Positions in which the body was aligned parallel to the nearest mirror (“parallel observations”) and at an angle to the mirror (“angled observations”: 1°-179° towards the left or right eye use) were scored jointly.

### Social Preference

Thirty larvae of the AB strain were examined at four weeks of age (Figure 2A), each experimental group was composed of 12 controls (CTRL), 11 and 7 larvae treated with 1μM VPA for 24 and 48 hours, respectively. Each group was observed only once. The apparatus consisted of a circular amaranth tank (diameter × height: 175 × 27 cm), surrounded by a circular black curtain fixed on a wood-and-metal frame in which the social preference apparatus was placed. The social preference apparatus was made of white plastic walls (11.2 × 4.2 × 8 cm) and divided into two chambers by a transparent barrier so that the experimental larva could see the companions. One of the chambers (the social chamber, 7 × 4.2 × 8 cm) hosted the experimental larva under test, while the other (4.2 × 4.2 × 8 cm) hosted 6 conspecifics (Figure 1A). To analyze the social preference, the social chamber was divided into two zones (3.5 × 4.2 × 8 cm), one proximal to the conspecifics and one distal from them. The water was 2.5 cm deep and was kept at 26 ± 1°C. The apparatus was lit from above (height: 30 cm) through a set of LED lights (∼ 1700 lumen). Each larva was placed in turn in the center of the social chamber and video-recorded using a high-resolution camera (FLIR Systems) for 5 min. The video recordings were coded offline, and the time spent by the larvae in the two zones was scored manually.

### Microdissection and RNA extraction

Three months-old AB zebrafish (Figure 3A and 4A; six controls and six treated at 5 hpf with 1μM VPA for 24 hrs) were anesthetized in an ice-cold water bath and sacrificed by decapitation; their brains were removed and dissected in ice-cold phosphate-buffered saline solution (PBS; Fisher Bioreagents, USA). The two hemispheres of telencephalon (including the pallium and the subpallium) and of the thalamus were collected separately from each animal and used for total RNA extractions using the RNeasy Mini Kit (QIAGEN; Milan, Italy). Briefly, tissues were homogenized in lysis buffer, run onto RNeasy spin columns, treated with DNase (RNase-Free DNase Set, QIAGEN; Milan, Italy) and eluted in RNase/DNase-free water. Total RNAs were quantified using NanoDrop™ (Thermo Fisher Scientific; Monza, Italy), and reverse transcribed using the SuperScript VILO™ cDNA Synthesis Kit (Invitrogen, Thermo Fisher Scientific; Monza, Italy) according to the manufacturer’s instructions.

### Quantitative real-time PCR

RT-qPCR experiments were performed using specific, commercially synthesized primer pairs (Merck Life Science Srl, Milan, Italy) as previously reported^59^. The triplicate reactions/samples were performed using the PowerUp™ SYBR™ Green Master Mix and a CFX96™ Real-Time System (Bio-Rad, Milan, Italy). The dCt method was used for expression quantification, raw expression data were normalized on the expression of the 18S reference gene. The lists of primers are reported in Table 5.

### Statistical analysis

In the mirror test, differential eye use was evaluated using the left visual field index, calculated as frequency of left eye use/(frequency of right eye use + frequency of left eye use). Values significantly higher than 0.5 indicate a preference for left eye use, while values significantly lower than 0.5 indicate a preference for right eye use. To evaluate the social preference of zebrafish larvae, the following variables were measured: the percentage [%] of total time spent (TTS) in the proximal zone (referred to as social preference index)^57^, the latency [s] to the first change of zone, and the number of alternations between the two zones (proximal-to-distal; distal-to-proximal) during the social preference test. Values of social preference index (%) were calculated as ((TTS in proximal zone/TTS proximal + TTS distal) x100), the values range from 100% (full choice for the social (proximal) zone) to 0% (full choice for the distal zone), where 50% represents the absence of preference. The effect of treatment on the left visual field index was evaluated by one-way analysis of variance (ANOVA), the effect of treatment and time on the social preference index, on the latency to first change of zone and on the spontaneous alternations was evaluated by multifactorial ANOVA. Statistical evaluation on the expression levels was performed on the log2 gene expression levels (dCt), the effect of treatment, brain side and transcript type was estimated using a linear mixed model, considering them as fixed factors and the experiment (experimental unit) as random factor. A Left Lateralization Index (LI) was calculated using the linear expression levels in the two hemispheres calculated as: LI = (left expression – right expression)/(left expression + right expression). For all the tests, significant departures of the social preference index/left visual field index/LI from chance level (50%, 0.5 and 0, respectively) were estimated by one-sample two-tailed t-tests. Graphs were generated with GraphPad Prism 9, statistical analyses were performed with Rstudio, using the *nlme*^1^ package for the linear mixed models and the *emmeans*^2^ package for Tukey pairwise comparison tests. Alpha was set to 0.05 for all tests.

### Data availability

The datasets generated and analysed during the current study are included in this published article as supplementary information files.

## Footnotes

1 https://cran.r-project.org/web/packages/nlme/index.html

2 https://cran.r-project.org/web/packages/emmeans/index.html

## Notes

### Competing Interest Statement

The authors have declared no competing interest.

### Summary of Updates

Result and Discussion revised to clarify significance of the results. Corresponding authorship updated.

