## Supplementary material for "Valproic acid exposure affects social visual lateralization and asymmetric gene expression in zebrafish larvae": Tables 1-4

| Table 1. Gene expression analysis in the thalamus: pairwise comparison report | | | | | | | | | | | | |
| --- | --- | --- | --- | --- | --- | --- | --- | --- | --- | --- | --- | --- |
| Thalamus | CTRL | | | | | | VPA | | | | | |
|  | ***dex*** | | ***lov*** | | ***ron*** | | ***dex*** | | ***lov*** | | ***ron*** | |
|  | **right** | **left** | **right** | **left** | **right** | **left** | **right** | **left** | **right** | **left** | **right** | **left** |
| Mean | 1.172 | 1.573 | -4.722 | -3.997 | 0.902 | 0.659 | 1.602 | 0.991 | -4.195 | -4.490 | 1.467 | 1.004 |
| Std. Error | 0.194 | 0.194 | 0.194 | 0.194 | 0.194 | 0.194 | 0.335 | 0.335 | 0.335 | 0.335 | 0.335 | 0.335 |
| N | 6 | 6 | 6 | 6 | 6 | 6 | 6 | 6 | 6 | 6 | 6 | 6 |
| Lower 95% C.L. | 0.674 | 1.075 | -5.220 | -4.495 | 0.404 | 0.162 | 0.740 | 0.129 | -5.057 | -5.352 | 0.605 | 0.142 |
| Upper 95% C.L. | 1.670 | 2.070 | -4.225 | -3.500 | 1.400 | 1.157 | 2.464 | 1.853 | -3.333 | -3.628 | 2.329 | 1.867 |
| Contrast (right - left) |  | |  | |  | |  | |  | |  | |
| t.ratio | -1.619 | | -2.930 | | 0.980 | | 2.470 | | 1.195 | | 1.871 | |
| p.value | 0.111 | | **0.005** | | 0.331 | | **0.017** | | 0.237 | | 0.067 | |
| η² (partial)^[1]^ | -0.935 | | -1.692 | | 0.566 | | 1.426 | | 0.690 | | 1.080 | |
| Lower 95% C.L. | -2.419 | | -3.176 | | -0.918 | | -0.058 | | -0.794 | | -0.404 | |
| Upper 95% C.I. | 0.549 | | -0.208 | | 2.050 | | 2.910 | | 2.174 | | 2.564 | |

| Table 2. Lateralisation Index in thalamus: one-sample t-test summary report | | | | | | |
| --- | --- | --- | --- | --- | --- | --- |
| Thalamus | CTRL | | | VPA | | |
|  | ***dex*** | ***lov*** | ***ron*** | ***dex*** | ***lov*** | ***ron*** |
| Mean | 0.136 | 0.246 | -0.076 | -0.200 | -0.098 | -0.153 |
| Std. Error | 0.061 | 0.021 | 0.108 | 0.086 | 0.079 | 0.097 |
| N | 6 | 6 | 6 | 6 | 6 | 6 |
| Lower 95% C.I. | -0.022 | 0.192 | -0.353 | -0.421 | -0.301 | -0.402 |
| Upper 95% C.I. | 0.293 | 0.299 | 0.200 | 0.021 | 0.106 | 0.095 |
| t.ratio | 2.216 | 11.734 | -0.708 | -2.326 | -1.235 | -1.587 |
| p.value | 0.078 | **<0.0001** | 0.510 | 0.068 | 0.272 | 0.173 |
| Cohen’s d^[2]^ | 0.90 | 4.79 | -0.29 | -0.95 | -0.50 | -0.65 |
| Lower 95% C.I. | -0.09 | 1.82 | -1.09 | -1.90 | -1.34 | -1.51 |
| Upper 95% C.I. | 1.84 | 7.77 | 0.54 | 0.06 | 0.37 | 0.27 |

| Table 3. Gene expression analysis in the telecephalon: pairwise comparison report | | | | | | | | | | | | |
| --- | --- | --- | --- | --- | --- | --- | --- | --- | --- | --- | --- | --- |
| Telencephalon | CTRL | | | | | | | | | | | |
|  | ***arbb2*** | | ***fez*** | | ***gap43*** | | ***nipa1*** | | ***nipa2*** | | ***robo1*** | |
|  | **right** | **left** | **right** | **left** | **right** | **left** | **right** | **left** | **right** | **left** | **right** | **left** |
| Mean | 2.966 | 3.047 | 3.013 | 3.009 | 4.565 | 4.763 | -2.083 | -2.118 | 0.182 | 0.138 | 1.452 | 1.688 |
| Std. Error | 0.283 | 0.283 | 0.283 | 0.283 | 0.283 | 0.283 | 0.283 | 0.283 | 0.283 | 0.283 | 0.283 | 0.283 |
| N | 4 | 4 | 4 | 4 | 4 | 4 | 4 | 4 | 4 | 4 | 4 | 4 |
| Lower 95% C.L. | 2.064 | 2.145 | 2.112 | 2.108 | 3.664 | 3.862 | -2.984 | -3.019 | -0.720 | -0.763 | 0.550 | 0.787 |
| Upper 95% C.L. | 3.867 | 3.948 | 3.915 | 3.911 | 5.466 | 5.665 | -1.181 | -1.216 | 1.083 | 1.040 | 2.353 | 2.590 |
| Contrast (right - left) |  | |  | |  | |  | |  | |  | |
| t.ratio | -0.400 | | 0.021 | | -0.981 | | 0.175 | | 0.214 | | -1.170 | |
| p.value | 0.691 | | 0.984 | | 0.330 | | 0.862 | | 0.831 | | 0.246 | |
| η² (partial)^1^ | -0.283 | | 0.015 | | -0.693 | | 0.124 | | 0.152 | | -0.827 | |
| Lower 95% C.L. | -2.533 | | -2.236 | | -2.944 | | -2.127 | | -2.099 | | -3.078 | |
| Upper 95% C.L. | 1.968 | | 2.265 | | 1.557 | | 2.374 | | 2.402 | | 1.423 | |
| Telencephalon | VPA | | | | | | | | | | | |
|  | ***arbb2*** | | ***fez*** | | ***gap43*** | | ***nipa1*** | | ***nipa2*** | | ***robo1*** | |
|  | **right** | **left** | **right** | **left** | **right** | **left** | **right** | **left** | **right** | **left** | **right** | **left** |
| Mean | 2.998 | 3.198 | 2.909 | 3.213 | 4.377 | 4.837 | -2.172 | -1.974 | -0.050 | 0.588 | 1.254 | 1.937 |
| Std. Error | 0.277 | 0.277 | 0.277 | 0.277 | 0.277 | 0.277 | 0.277 | 0.277 | 0.277 | 0.277 | 0.277 | 0.277 |
| N | 4 | 4 | 4 | 4 | 4 | 4 | 4 | 4 | 4 | 4 | 4 | 4 |
| Lower 95% C.L. | 2.116 | 2.317 | 2.028 | 2.332 | 3.496 | 3.956 | -3.053 | -2.855 | -0.931 | -0.294 | 0.373 | 1.056 |
| Upper 95% C.L. | 3.879 | 4.079 | 3.790 | 4.094 | 5.258 | 5.719 | -1.291 | -1.093 | 0.831 | 1.469 | 2.135 | 2.818 |
| Contrast (right - left) |  | |  | |  | |  | |  | |  | |
| t.ratio | -0.993 | | -1.504 | | -2.279 | | -0.977 | | -3.152 | | -3.377 | |
| p.value | 0.324 | | 0.137 | | **0.026** | | 0.332 | | **0.002** | | **0.001** | |
| η² (partial)^1^ | -0.702 | | -1.063 | | -1.611 | | -0.690 | | -2.229 | | -2.388 | |
| Lower 95% C.L. | -2.952 | | -3.314 | | -3.861 | | -2.941 | | -4.479 | | -4.638 | |
| Upper 95% C.L. | 1.548 | | 1.187 | | 0.639 | | 1.560 | | 0.022 | | -0.137 | |

| Table 4. Lateralisation Index in telencephalon: one-sample t-test summary report | | | | | | | | | | | | |
| --- | --- | --- | --- | --- | --- | --- | --- | --- | --- | --- | --- | --- |
| Telencephalon | CTRL | | | | | | VPA | | | | | |
|  | ***arbb2*** | ***fez*** | ***gap43*** | ***nipa1*** | ***nipa2*** | ***robo1*** | ***arbb2*** | ***fez*** | ***gap43*** | ***nipa1*** | ***nipa2*** | ***robo1*** |
| Mean | 0.028 | -0.001 | 0.067 | -0.012 | -0.015 | 0.080 | 0.069 | 0.105 | 0.157 | 0.068 | 0.217 | 0.228 |
| Std. Error | 0.047 | 0.021 | 0.079 | 0.050 | 0.077 | 0.089 | 0.013 | 0.028 | 0.050 | 0.043 | 0.037 | 0.079 |
| N | 4 | 4 | 4 | 4 | 4 | 4 | 4 | 4 | 4 | 4 | 4 | 4 |
| Lower 95% C.I. | -0.123 | -0.067 | -0.184 | -0.171 | -0.261 | -0.203 | 0.028 | 0.015 | -0.003 | -0.068 | 0.098 | -0.022 |
| Upper 95% C.I. | 0.179 | 0.064 | 0.319 | 0.146 | 0.231 | 0.363 | 0.111 | 0.194 | 0.318 | 0.204 | 0.335 | 0.478 |
| t.ratio | 0.586 | -0.070 | 0.851 | -0.246 | -0.191 | 0.899 | 5.336 | 3.728 | 3.116 | 1.596 | 5.818 | 2.904 |
| p.value | 0.599 | 0.948 | 0.457 | 0.821 | 0.861 | 0.435 | **0.013** | **0.034** | 0.053 | 0.209 | **0.010** | 0.062 |
| Cohen’s d^2^ | 0.29 | -0.04 | 0.43 | -0.12 | -0.10 | 0.45 | 2.67 | 1.86 | 1.56 | 0.80 | 2.91 | 1.45 |
| Lower 95% C.I. | -0.73 | -1.01 | -0.64 | -1.10 | -1.07 | -0.62 | 0.42 | 0.12 | -0.01 | -0.39 | 0.51 | -0.06 |
| Upper 95% C.I. | 1.28 | 0.95 | 1.43 | 0.87 | 0.89 | 1.46 | 4.90 | 3.55 | 3.06 | 1.91 | 5.31 | 2.89 |

1. Effect size calculated using the R package *emmeans*
2. Effect size calculated using the R package *effectsize*
