## Supplementary material for "Valproic acid exposure affects social visual lateralization and asymmetric gene expression in zebrafish larvae": Table 5

| Table 5. List of primers used in the study | | | | | |
| --- | --- | --- | --- | --- | --- |
| Gene name | Gene ID | Primers | Sequence (5’- 3’) | Amplicon size | Efficiency (%) |
| ***18S ribosomal RNA*** | 100037361 | For | TCGCTAGTTGGCATCGTTTATG | 85 bp | 93 |
|  |  | Rev | CGGAGGTTCGAAGACGATCA |  |  |
| ***kctd12.1/lov*** | 373865 | For | AGTTCTTTCAGCTGCGGGACCTTA | 112 bp | 98.3 |
|  |  | Rev | GCGACGGACAGTGTGCGAGAG |  |  |
| ***kctd8/dex*** | 568933 | For | CATGCCATCAATTACGCAAC | 112 bp | 101.9 |
|  |  | Rev | CGGATCCCGACTTTCATTTA |  |  |
| ***kctd12.2/ron*** | 553403 | For | GCCACTCAACTTTGCTCTCC | 191 bp | 99.7 |
|  |  | Rev | GAGGCTCGCTTTCTCTTTCTCTTTGA |  |  |
| ***arrb2*** | 394099 | For | CCCTTCAACTTCACGATTCC | 127 bp | 93 |
|  |  | Rev | CATCCACAGTTTTGGCACAG |  |  |
| ***fez1*** | 406705 | For | AAACACTCAACGGCAACCTC | 159 bp | 95.2 |
|  |  | Rev | CCGGTGAGTTTTCCATCATC |  |  |
| ***gap43*** | 30608 | For | GACACATCACCCGGAAAAAG | 151 bp | 96.4 |
|  |  | Rev | TCATTTGCTGGGGAGTTAGG |  |  |
| ***robo1*** | 30769 | For | ATCTCAATCCCGAAGTGCTG | 118 bp | 97.5 |
|  |  | Rev | TTTCCTGCTCACAGACGATG |  |  |
| ***nipa1*** | 450042 | For | AGTCTTGTTTGGTGCCGTTC | 119 bp | 101.4 |
|  |  | Rev | TGGGTGAGTGAATGATGAGC |  |  |
| ***nipa2*** | 406399 | For | CTTTGTGGTGTTTGCGACAG | 176 bp | 98.5 |
|  |  | Rev | CAGCGATGGCCTCTTTAATC |  |  |
